# Hippocampal gamma-band oscillopathy in a mouse model of Fragile X Syndrome

**DOI:** 10.1101/2021.04.24.441239

**Authors:** Evangelia Pollali, Jan-Oliver Hollnagel, Gürsel Çalışkan

## Abstract

Fragile X syndrome (FXS) is the most common inherited form of intellectual disability arising from the loss of fragile X mental retardation protein (FMRP), a protein that plays a central role in neuronal function and plasticity. FXS patients show sensory hypersensitivity, hyperarousal and hippocampus-dependent learning deficits that can be recapitulated in the FMR1 KO mice. Enhanced metabotropic glutamate receptor (mGluR) signaling and muscarinic acetylcholine receptor (mAChR) signaling in the FMR1 KO mouse are implicated as the primary causes of the disease pathogenesis. Furthermore, glutamatergic kainate receptor (KAR) function is reduced in the cortex of the FMR1 KO mice. Of note, activation of these signaling pathways leads to slow gamma-range oscillations in the hippocampus *in vitro* and abnormal gamma oscillations have been reported in FMR1 KO mice and patients with FXS. Thus, we hypothesized that aberrant activation of these receptors leads to the observed gamma oscillopathy. We recorded gamma oscillations induced by either cholinergic agonist carbachol (CCh), mGluR1/5 agonist Dihydroxyphenylglycine (DHPG) or ionotropic glutamatergic agonist KA from the hippocampal CA3 in WT and FMR1 KO mice *in vitro*. We show a specific increase in the power of DHPG and CCh-induced gamma oscillations and reduction in the synchronicity of gamma oscillations induced by KA. We further elucidate an aberrant spiking activity during CCh-induced and kainate-induced gamma oscillations which may underlie the altered gamma oscillation synchronization in the FMR1 KO mice. Last, we also noted a reduced incidence of spontaneously-occurring hippocampal sharp wave-ripple events. Our study provides further evidence for aberrant hippocampal rhythms in the FMR1 KO mice and identifies potential signaling pathways underlying gamma band oscillopathy in FXS.

## 1. Introduction

Expansion of non-coding trinucleotide CGG above the normal range (>200 CGG repeats) causes the full mutation and loss of fragile X mental retardation protein (FMRP) leading to Fragile X syndrome (FXS), the most common genetically inherited form of intellectual disability and autism spectrum disorder (Hagerman et al., 2017; Salcedo-Arellano et al., 2020). Lack of FMRP, a key translational regulator of vast majority of synaptic proteins, leads to excessive protein synthesis, impaired synaptic function and dysregulation of protein-synthesis dependent plasticity (Bassell and Warren, 2008; Santos et al., 2014). Accumulating evidence suggests that elevated muscarinic acetylcholine receptor (mAChR)-mediated signaling and metabotropic glutamate receptor5 (mGluR5)-mediated signaling are involved in the pathogenesis of FXS (Bear et al., 2004; Dölen et al., 2007; Thomson et al., 2017). It is widely-accepted that loss of FMRP leads to excessive mGluR5-mediated protein synthesis and enhanced mGluR-mediated long-term depression (LTD) (Bear et al., 2004; Dölen and Bear, 2008; Huber et al., 2002). Importantly, both electrophysiological and behavioral phenotypes observed in the FMR1 KO mice can be corrected by genetic or pharmacological reduction of mGluR5 signaling (Dölen et al., 2007; Michalon et al., 2012). Similarly, FMR1 KO mice show increased expression of both type 1 (M1) and type 4 (M4) mAChR in the hippocampus (Thomson et al., 2017). Accordingly, hippocampal LTD induced by activation of mAChR is exaggerated (Volk et al., 2007) and antagonism of M1 can rescue some of the behavioral phenotypes observed in FMR1 KO mice (Veeraragavan et al., 2011). On the other hand, ionotropic glutamate receptor function is also affected by lack of FMRP (Aloisi et al., 2017; Pilpel et al., 2009). Interestingly, a reduced kainate receptor (KAR)-mediated synaptic function has only recently been demonstrated in the cortex of the FMR1 KO mice (Qiu et al., 2018).

Network oscillations are electrical brain activities reflecting neuronal synchronization at different spatial and temporal levels (Buzsaki and Draguhn, 2004). Neuronal network oscillations are crucial for maintaining most of the brain functions and can be used as proxies for neural mechanisms underlying behaviorally-relevant synaptic plasticity (Çalışkan and Stork, 2018). While the molecular and cellular mechanisms underlying FXS have been extensively studied in rodent models, systematic investigation of behaviorally-relevant neurophysiological oscillations as mesoscopic biomarkers for FXS has only recently accelerated (Arbab et al., 2018; Boone et al., 2018; Dvorak et al., 2018; Jonak et al., 2020; Lovelace et al., 2020, 2018; Pirbhoy et al., 2020; Radwan et al., 2016; Wen et al., 2019; Wong et al., 2020). Of note, the most striking and converging finding from studies using FMR1 KO mouse model is the gamma-range (30-80 Hz) oscillopathy observed in distinct cortical regions (Jonak et al., 2020; Lovelace et al., 2020, 2018; Pirbhoy et al., 2020; Wen et al., 2019; Wong et al., 2020) and hippocampus (Arbab et al., 2018; Boone et al., 2018; Dvorak et al., 2018; Radwan et al., 2016). Elevated gamma oscillations appear to be present during diverse behavioral stages including active exploration, rest and sleep (Arbab et al., 2018; Boone et al., 2018; Lovelace et al., 2018). Similar enhancement in the resting state gamma power and aberrant synchronization of gamma oscillations have been reported in electroencephalogram (EEG) studies in FXS patients (Ethridge et al., 2019, 2017; Wang et al., 2017). These oscillatory changes have been associated with sensory and cognitive deficits in FXS (Sinclair et al., 2017). Specifically, enhanced hippocampal gamma oscillations and their aberrant synchronization have been implicated in reduced cognitive flexibility in the FMR1 KO mice (Dvorak et al., 2018; Radwan et al., 2016). Thus, aberrant gamma oscillations might indeed be a common mesoscopic biomarker for FXS and potentially for autism spectrum disorders. However, to date, it is still not clarified whether elevated mGluR signaling and/or mAChR signaling can indeed lead to an aberrant enhancement of gamma oscillations in the FMR1 KO mice.

*In vitro* oscillation models provide a robust tool for pharmacological studies aiming at elucidating phenotypic differences (Albrecht et al., 2013; Hollnagel et al., 2019; Klein et al., 2016). Accumulating evidence suggests that *in vitro* network oscillations engage similar cellular interactions as observed *in vivo* (Butler and Paulsen, 2015; Buzsáki, 2015; Schlingloff et al., 2014) and are sensitive to genetic and/or behavioral manipulations (Albrecht et al., 2013; Caliskan et al., 2015; Çalışkan et al., 2016; Hollnagel et al., 2019; Lu et al., 2011; Mizunuma et al., 2014; Norimoto et al., 2018). Specifically, the hippocampal CA3 subregion with its tightly interconnected recurrent network can generate gamma-range rhythmic local field potentials (LFP) *in vitro* (Fisahn et al., 1998; Le Duigou et al., 2014). These LFP oscillations can be induced via activation of mAChR (Caliskan et al., 2015; Fisahn et al., 2002; Wójtowicz et al., 2009; Zemankovics et al., 2013) or elevating glutamatergic input via activation of either mGluR1/5 (Gillies et al., 2002; Pálhalmi et al., 2004) or KAR (Albrecht et al., 2013; Fisahn et al., 2004; Wójtowicz et al., 2009; Zemankovics et al., 2013). In the absence of elevated cholinergic or glutamatergic input, hippocampal CA3 network generates spontaneous sharp wave-ripple (SW-R) activity *in vitro* that represents a distinct hippocampal network state (Buzsáki, 2015; Maier et al., 2003). Thus, two distinct hippocampal LFP states observed *in vivo* can partially be mimicked using hippocampal slice preparations.

In the current study, we recorded *in vitro* hippocampal network oscillations in the FMR1 KO mice and analyzed spontaneous SW-Rs and pharmacologically-induced gamma oscillations via activating mAChR, mGluR1/5 or KAR. Based on the elevated mAChR- and mGluR5-mediated signaling in the FMR1 KO mice, we postulated that direct activation of these signaling pathways *in vitro* would be sufficient to observe the aberrantly elevated gamma oscillations in FMR1 KO mice. Our findings provide promising evidence for direct involvement of elevated mAChR and mGluR5-mediated signaling in the gamma-range oscillopathy observed in FXS.

## 2. Materials and methods

### 2.1. Animals

Male FMR1 KO (−/y) mice and wild-type (WT) littermates (+/y) with FVB background (JAX stock #003024)(Bakker et al., 1994) were bred in our animal facility, at the Institute of Biology, Otto-von-Guericke University Magdeburg (12h light/dark cycle with lights switched on at 7 p.m. with a 30 min dawn phase; food and water ad libitum). All experiments were conducted in accordance with the European and German regulations for animal experiments and were approved by the Landesverwaltungsamt Saxony-Anhalt (Permission No. 42502-2-1009-UniMD).

### 2.2. Electrophysiology

#### 2.2.1. Slice Preparation

Animals were anaesthetized with isoflurane and decapitated. The brain was rapidly removed and submerged in ice-cold artificial cerebrospinal fluid (aCSF) including (in mM) 129 NaCl, 21 NaHCO_3_, 3 KCl, 1.6 CaCl_2_, 1.8 MgCl, 1.25 NaH_2_PO4 and 10 glucose (carbogenated with 5% CO_2_, 95% O_2_, pH 7.4, osmolarity 300 mOsm). Horizontal slices containing ventral-to-mid hippocampus (400 μm) were prepared with a vibratome (Model 752; Campden Instruments LTD, Loughborough, UK) at an angle of about 12° in the fronto-occipital direction and placed in an interface chamber at 32 °C (aCSF flow rate: ~2 ml/min). Slices were allowed to recover for at least 1 h before starting the recordings.

#### 2.2.2. Local field potentials

LFP recordings were obtained from stratum pyramidale (SP) of CA3 and CA1. Borosilicate glass electrodes of ~1 MΩ resistance, filled with aCSF, were placed at a depth of ~80 μm and spontaneous SW-R were recorded. Gamma oscillations were induced by increasing the temperature to 35 °C and bath-application of drugs at least for 45 min. The signal was amplified by a commercial (EXT-02F – Extracellular Amplifier; npi Electronic GmbH, Tamm, Germany) or a custom-made amplifier and low-pass filtered at 3 kHz. The data were sampled at a rate of 5 kHz using a data acquisition unit (CED-Mikro1401; Cambridge Electronic Design, Cambridge, UK). Signals were recorded using Spike2.8 software (Cambridge Electronic Design) and stored on a computer hard disc and analyzed off-line.

### 2.3. Drug Application

Drugs were applied via continuous perfusion with a flow rate of ~2 ml/min. AChR agonist Carbachol (CCh; 5 μM) (TOCRIS, Bristol, UK), mGluR1/5 agonist Dihydroxyphenylglycine (DHPG; 10 μM) (TOCRIS) and ionotropic GluR agonist Kainic acid (KA; 60 nM) (TOCRIS) were added to aCSF freshly prior to induction of gamma oscillations, using stock aliquots stored in −20°C. CCh stocks were aliquoted in dimethyl sulfoxide (DMSO; Final Concentration: 0.01 %) (Merck KGaA, Darmstadt, Germany) whereas DHPG and KA were aliquoted in distilled water.

### 2.4. Data analysis

#### 2.4.1. Gamma oscillation analysis

Pharmacologically-induced gamma oscillations were recorded from pyramidal layer of CA3 region of ventral-to-mid hippocampus (Fig. 1). Two minutes artefact-free data were extracted and analyzed by Fast Fourier Transformation using Spike2.8 software (Cambridge Electronic Design) with a frequency resolution of 1.221 Hz and custom-made MATLAB scripts (MathWorks, Natick, MA, USA). Peak frequency, integrated power (20-80 Hz) and half band width (HBW) were calculated from the power spectra of two-minute artifact free data. The slices that show gamma oscillations with peak power greater than 10 μV^2^ and peak frequency greater than 20 Hz were included in the analysis. HBW was calculated as the frequency range at the half-maximum of the peak power. Autocorrelation was calculated as the 2^nd^ positive peak of two-minute auto-correlogram and time constant (Tau) was calculated as the decaying exponential fit to the peaks of the autocorrelation.

**Fig. 1.**
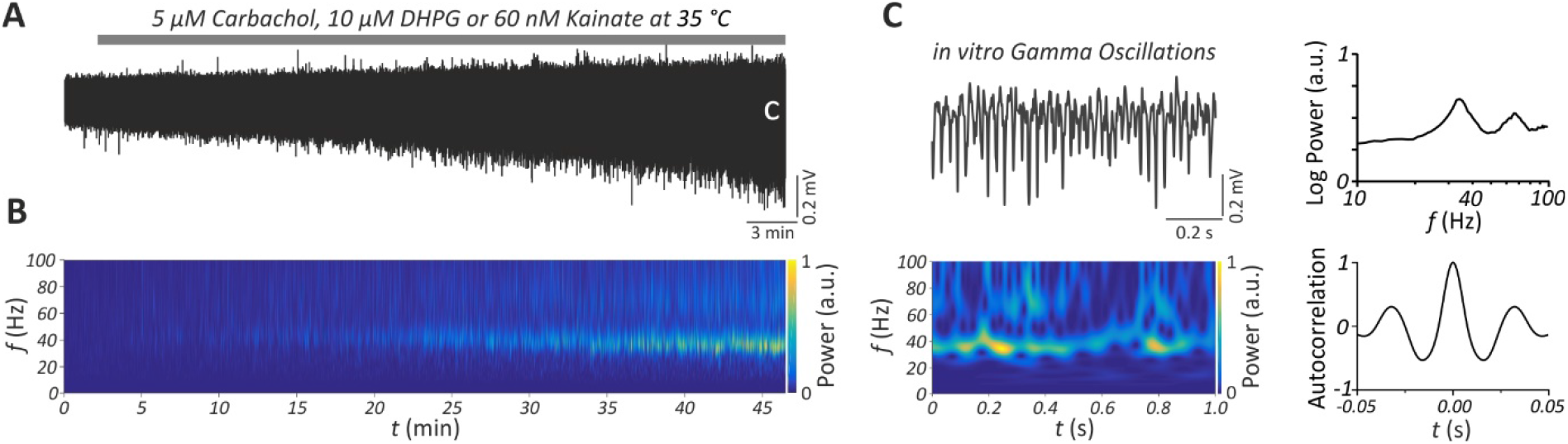
Gamma oscillations in the hippocampal CA3 *in vitro*. **(A)** Sample trace of gamma oscillations induced via bath-application of either Carbachol (CCh, 5 μM), DHPG (10 μM) or Kainate (KA, 60 nM) for 45-60 min. Local field potential (LFP) recordings were performed from the pyramidal layer of CA3 subregion in horizontal brain slices including transverse-like sections of the ventral-to-mid hippocampus. “C” indicates the gamma oscillation trace shown in panel C. **(B)** Corresponding wavelet transformation of the LFP trace in panel A showing power of frequency domains (*f*) over time (*t*). Heat scale colors encode for Power in arbitrary units (a.u.). Note the emergence of gamma oscillations at 30-40 Hz after 15 min of drug application and its stabilization after ~45 min. **(C)** Sample LFP gamma trace (left, up) and corresponding wavelet transformation (left, down) ~45 min after drug perfusion. Note the prominent gamma oscillations at ~40 Hz. Power Spectra (right, up) or auto-correlograms (right, down) were generated from artefact-free two min gamma oscillation segments at least 45 min after perfusion of CCh, DHPG or KA. Note that the *f* is shown in logarithmic scale.

#### 2.4.2. Multi-unit activity analysis

Multi-unit activity during gamma oscillations was analyzed adapting previously described methods (Hájos et al., 2004). In brief, gamma oscillations were band-pass filtered at 15–45 Hz (FFT filter) and further processed by a Hilbert transformation yielding the phase representation of gamma cycles. Unit activity was detected by extracting the high frequency component of the LFP signal (FFT filter: 500– 3000 Hz). The band-pass filtered signal was then denoised by application of the wavelet denoising function implemented in MATLAB’s Wavelet Toolbox. For initial collection of units, only those with amplitudes bigger than 0.0125 mV were considered. Further evaluation was done with units representing the upper quartile of unit amplitudes. Unit timings were then related to the phase of the corresponding gamma cycle. For comparison of units and respective gamma cycles, gamma cycles and units’ spike timings were resampled. For each recording, the obtained gamma cycles were averaged separately. The polar distribution of units was normalized to represent the probability of unit firing within a gamma cycle. For statistical evaluation, the unit firing probability curves and their activation curves (i.e. cumulative firing probability) were plotted in two dimensions. Of note, we compared multi-unit activity only in those experiments in which the gamma oscillations were clearly recorded in the pyramidal cell layer and thus providing representative activation curves.

#### 2.4.3. Sharp wave-ripple analysis

Extracellular local field potential recordings in the CA3 and CA1 subregions of the ventral-to-mid hippocampus exhibit spontaneous sharp wave-ripple (SW-R) activity *in vitro* (Çalışkan et al., 2016; Maier et al., 2003). Two minutes artefact-free data were extracted and custom-made MATLAB scripts (MathWorks, Natick, MA, USA) were used to analyze the sharp wave (SW) incidence, SW area under curve, the ripple frequency and ripple amplitude. Specifically, SW were identified by low-pass filtering the raw data at 45 Hz (FFT filter, cut frequency: 45 Hz). Only events greater than 3 times the standard deviation (SD) of the low-pass-filtered signal and with a minimum distance of 100 ms between two consecutive SW were considered for analysis. SW area was calculated by using as start and end of a SW event, the points crossing the mean of the data. In order to isolate the ripple component, time windows of 125 ms centered to the maximum of SW event were kept and band-pass filtered at 120– 300 Hz (FFT filter). Only ripples with amplitude greater than 3 times the SD of the band-pass-filtered signal were considered. Data with length of 15 ms before and 10 ms after the maximum of SW event were stored and ripple amplitude was calculated using triple-point-minimax-determination. Ripples were included in the analysis, when the difference between the falling and rising component of a ripple was lower than 75%. Ripple frequencies were calculated using the duration between the through of consecutive ripples within each SW-R complex.

### 2.5. Statistical analysis

The number of animals (N) and the number of slices (n) used are indicated in the figure legends. GraphPad Prism version 9.0.0 for Windows (GraphPad Software, CA, USA) was used for statistical comparison. First, Shapiro–Wilk test was used to determine whether the data are normally distributed. The genotype differences were determined either by Student’s two-tailed t-test for normally-distributed data and Mann-Whitney U test was used otherwise. Log transformation was used for the statistical comparison of gamma power due to the variable nature of gamma oscillation power *in vitro*. To provide information about differences in the multi-unit activity relative to the gamma cycle multiple t-test comparison was applied without assuming a similar standard deviation. Data are reported as mean ± standard error of the mean (SEM). Probability value p<0.05 was considered significant.

## 3. Results

### 3.1. Cholinergic gamma oscillations are enhanced in the CA3 of FMR1 KO mice

Converging evidence indicates that mAChR signaling is elevated in the FMR1 KO mice (Thomson et al., 2017; Veeraragavan et al., 2011; Volk et al., 2007). Furthermore, induction of cholinergic-type gamma oscillations depends on activation of mAChR (Fisahn et al., 2002, 1998). Thus, we first characterized cholinergic gamma oscillations in the hippocampal CA3 subregion of FMR1 KO and WT mice via bath-application of 5 μM CCh. Gamma oscillations emerged 10-15 min after application of CCh and stabilized after ~45 min (Fig. 2A). Strikingly, we found a profoundly increased gamma power (20-80 Hz) in the FMR1 KO compared to the WT mice (Fig. 2B, D, I; Student’s two-tailed t-test; T(57) = 4.055, p < 0.05). However, analysis of the gamma peak frequency (Fig. 2E; Mann-Whitney U test, p = 0.1463), the half band width (Fig. 2F; Mann-Whitney U test, p = 0.7477), the decay constant gamma autocorrelation fit (Tau) (Fig. 2G; Mann-Whitney U test, p = 0.5094) and the autocorrelation (2^nd^ peak value) of local gamma oscillations (Fig. 2C, H; Mann-Whitney U test; p = 0.4535) revealed no genotype differences. Of note, we observed significant differences in the unit activity which corresponds to an overall neuronal activity most likely reflecting action potentials generated by pyramidal cells (Fig. 2I-J). While the probability of unit firing was slightly higher at the beginning of each gamma cycle in WT mice, cells were more active in the late phase of gamma cycles in FMR1 KO mice (Fig. 2K-L; multiple t-test comparisons; p < 0.05, indicated by colored patches). Together, these data suggest that cholinergic gamma oscillations in the hippocampus of FMR1 KO mice are substantially augmented and may underlie the aberrantly enhanced gamma oscillations observed in FMR1 KO mice *in vivo*.

**Fig. 2.**
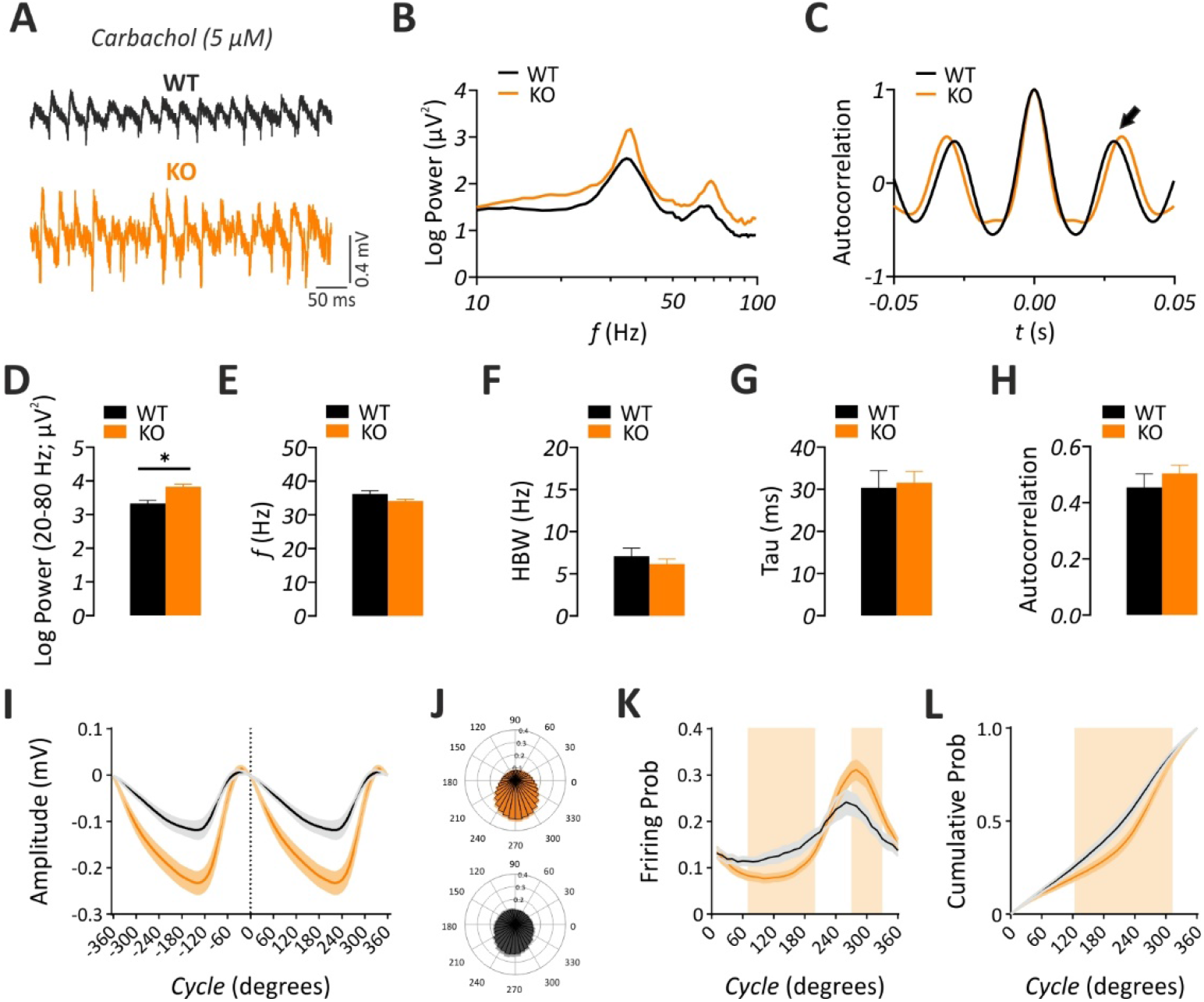
Carbachol-induced cholinergic gamma oscillations are augmented in the hippocampal CA3 subregion of FMR1 KO mice *in vitro*. **(A)** Sample trace of gamma oscillations induced via bath-application of Carbachol (CCh, 5 μM). **(B)** Corresponding power spectra of gamma oscillations showing a profound increase in the power of gamma-range (20-80 Hz) LFP oscillations without any shift in the gamma peak frequency (*f*). Note that the *f* is shown in logarithmic scale. **(C)** Representative autocorrelogram showing no change between genotypes. 2^nd^ positive peak value (indicated by the arrow) was measured for statistical comparison. Summary plots highlighting **(D)** a significant increase in the gamma power (WT: n = 24 / N = 8, KO: n = 35 / N = 11) and no significant alterations in **(E)** the gamma peak *f* (WT: n = 24 / N = 8, KO: n = 35 / N = 11), **(F)** half band width (HBW) (WT: n = 18 / N = 8, KO: n = 30 / N = 11), **(G)** time constant (Tau) of the decaying exponential fit to the peaks of the autocorrelation (WT: n = 18 / N = 8, KO: n = 30 / N = 11) and **(H)** 2^nd^ positive peak value of the autocorrelogram (WT: n = 18 / N = 8, KO: n = 30 / N = 11). Evaluation of multi-unit activity in datasets (WT: n = 12 / N = 6, KO: n = 25 / N = 8) is shown in **I-L**. **(I)**Average gamma cycle. Note that the cycle’s amplitude is plotted against degrees. **(J)** Polar plots illustrating the probability of unit activity in FMR1 KO (top) and WT (bottom) mice relative to the cycle shown in **I**. **(K)** Unit firing probability reveals different activation regimes. Units in WT are significantly more active in the initial phase, whereas units in FMR1 KO mice have a significantly higher firing probability in the late phase. **(L)** Unit activation curves reveal more uniform recruitment of units in WT mice. Statistical comparison was performed using Student’s two-tailed t-test for **C** and Mann-Whitney U test for **D**, **E**, **F** and **G**. In **K** and **L** statistical comparison was done with multiple t-test comparisons. Colored patches indicate ranges in which datasets differed significantly. Data are given as mean ± SEM. *p < 0.05.

### 3.2. DHPG-induced gamma oscillations are enhanced in the CA3 of FMR1 KO mice

In addition to mAChR signaling, increased mGluR signaling is also evident in the FMR1 KO mice (Dölen et al., 2007; Michalon et al., 2012). Activation of mGluR1/5 signaling in the hippocampus *in vitro* also induces gamma-range LFP oscillations (Gillies et al., 2002; Pálhalmi et al., 2004). Thus, we characterized the gamma network activity induced by activation of mGluR1/5 via DHPG (Fig. 3A). We found that the power of DHPG-induced gamma oscillations was significantly increased in the FMR1 KO compared to WT (Fig. 3B, D, I; Student’s two-tailed t-test; T(46) = 2.179, p < 0.05). As reported previously (Pálhalmi et al., 2004), DHPG-induced gamma oscillations showed an increased gamma peak frequency in comparison to the CCh-induced gamma oscillations (DHPG: 44.4 ± 1.5 Hz vs. CCh: 34 ± 0.2 Hz). However, no genotype differences were detected for gamma peak frequency (Fig. 3E; Student’s two-tailed t-test; T(46) = 0.8122, p = 0.4209). Similarly, HBW (FIG. 3F; Mann-Whitney U test, p = 0.4876), Tau (FIG. 3G; Mann-Whitney U test, p = 0.7788) and gamma autocorrelation (FIG. 3C, H; Student’s two-tailed t-test; T(30) = 0.6405, p = 0.5267) were comparable between FMR1 KO and WT mice. Thus, in addition to the enhanced cholinergic signaling, mGluR signaling might also be directly involved in the enhancement of gamma oscillations in the FMR1 KO mice. Interestingly, the evaluation of the multi-unit activity did not reveal an apparent difference between WT and FMR1 KO mice in the recruitment of putative pyramidal cells to generate action potentials during gamma cycles (Fig. 3I-L; multiple t-test comparisons; p > 0.05, except for one data point in Fig. 3K where p < 0.05).

**Fig. 3.**
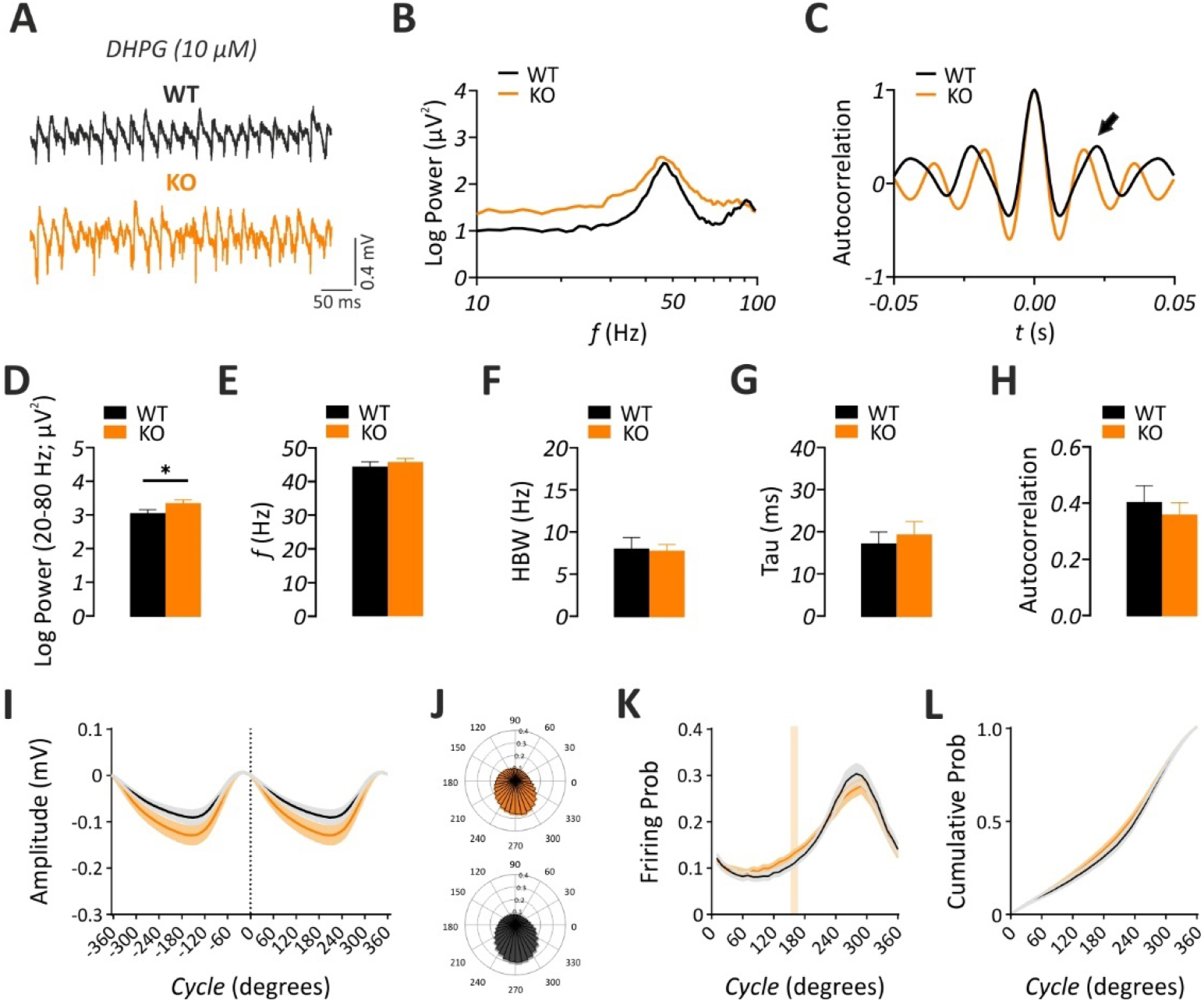
DHPG-induced gamma oscillations are augmented in the hippocampal CA3 subregion of FMR1 KO mice *in vitro*. **(A)** Sample trace of gamma oscillations induced via bath-application of Dihydroxyphenylglycine (DHPG, 10 μM). **(B)** Corresponding power spectra of gamma oscillations showing an increase in the power of gamma-range oscillations without any shift in the gamma peak oscillation frequency (*f*). Note that the *f* is shown in logarithmic scale. **(C)** Representative autocorrelogram showing no change between genotypes. 2^nd^ positive peak value (indicated by the arrow) was measured for statistical comparison. Summary plots illustrating a significant increase in **(D)** the gamma power (20-80 Hz) (WT: n = 22 / N = 9, KO: n = 26 / N = 8) and no significant alteration **(E)** in the gamma peak *f* (WT: n = 22 / N = 9, KO: n = 26 / N = 8), **(F)** half band width (HBW) (WT: n = 14 / N = 7, KO: n = 18 / N = 7), **(G)** time constant (Tau) of the decaying exponential fit to the peaks of the autocorrelation (WT: n = 14 / N = 7, KO: n = 18 / N = 7) and **(H)** 2^nd^ peak value of the autocorrelogram (WT: n = 14 / N = 7, KO: n = 18 / N = 7). Evaluation of multi-unit activity in datasets (WT: n = 20 / N = 8, KO: n = 24 / N = 8) is shown in **I-L**. **(I)** Average gamma cycle. Note that the cycle’s amplitude is plotted against degrees. **(J)** Polar plots illustrating the probability of unit activity in FMR1 KO (top) and WT (bottom) mice relative to the cycle shown in **I**. **(K)** Unit firing probability showing similar activation regimes. **(L)** Unit activation curves with similar recruitment of units in WT vs. FMR1 KO mice. Statistical comparison was performed using Student’s two-tailed t-test for **C, D, G** and Mann-Whitney U test for **E** and **F**. In **K** and **L** statistical comparison was done with multiple t-test comparisons. Colored line in **K** indicates significant difference. Data are given as mean ± SEM. *p < 0.05.

### 3.3. Synchronization of KA-induced gamma oscillations is reduced in the hippocampal CA3 of FMR1 KO mice

KAR activation has been a standard protocol for investigation of gamma oscillations *in vitro* (Bartos et al., 2007). In comparison to mAChR and mGluR signaling, a possible alteration in the ionotropic KAR signaling in the pathogenesis of FXS has been inadequately addressed. Thus, we aimed to characterize KA-induced gamma oscillations in the hippocampal CA3 of FMR1 KO mice (Fig. 4A). Interestingly, we found no alteration in the power of KA-induced gamma oscillations (Fig. 4B, D, I; Mann-Whitney U test, p =0.0959). On the other hand, the peak frequency of gamma oscillations was significantly increased (Fig. 4B, E; Student’s two-tailed t-test; T(59) = 2.009, p < 0.05). A tendency for an increased HBW (Fig. 4F; Mann-Whitney U test; p = 0.0598) was evident suggesting a possible change in the gamma synchronicity. Accordingly, analysis of time constant (Tau) of the decaying exponential fit to the peaks of the autocorrelation revealed a significant reduction (Fig. 4G; Student’s two-tailed t-test; T(50) = 3.056, p < 0.05) and a tendency for decreased autocorrelation (FIG. 4C, H; Student’s two-tailed t-test; T(50) = 2.001, p = 0.0508) in FMR1 KO mice compared to WT. At the multi-unit level and opposed to the findings described for CCh-induced gamma oscillations, we detected a more uniform activation curve in the in FMR1 KO mice (Fig 4I-L; multiple t-test comparisons; p < 0.05, indicated by colored patches). Here, the probability of unit firing was slightly higher at the beginning of the gamma cycle in FMR1 KO mice, while it was lower towards the end of the gamma cycle (Fig 4K; multiple t-test comparisons; p < 0.05, indicated by colored patches). These data suggest a general reduction in the synchronicity of KA-induced gamma oscillations and aligns with a reduction KAR-mediated synaptic function reported in the cortex of FMR1 KO mice (Qiu et al., 2018).

**Fig. 4.**
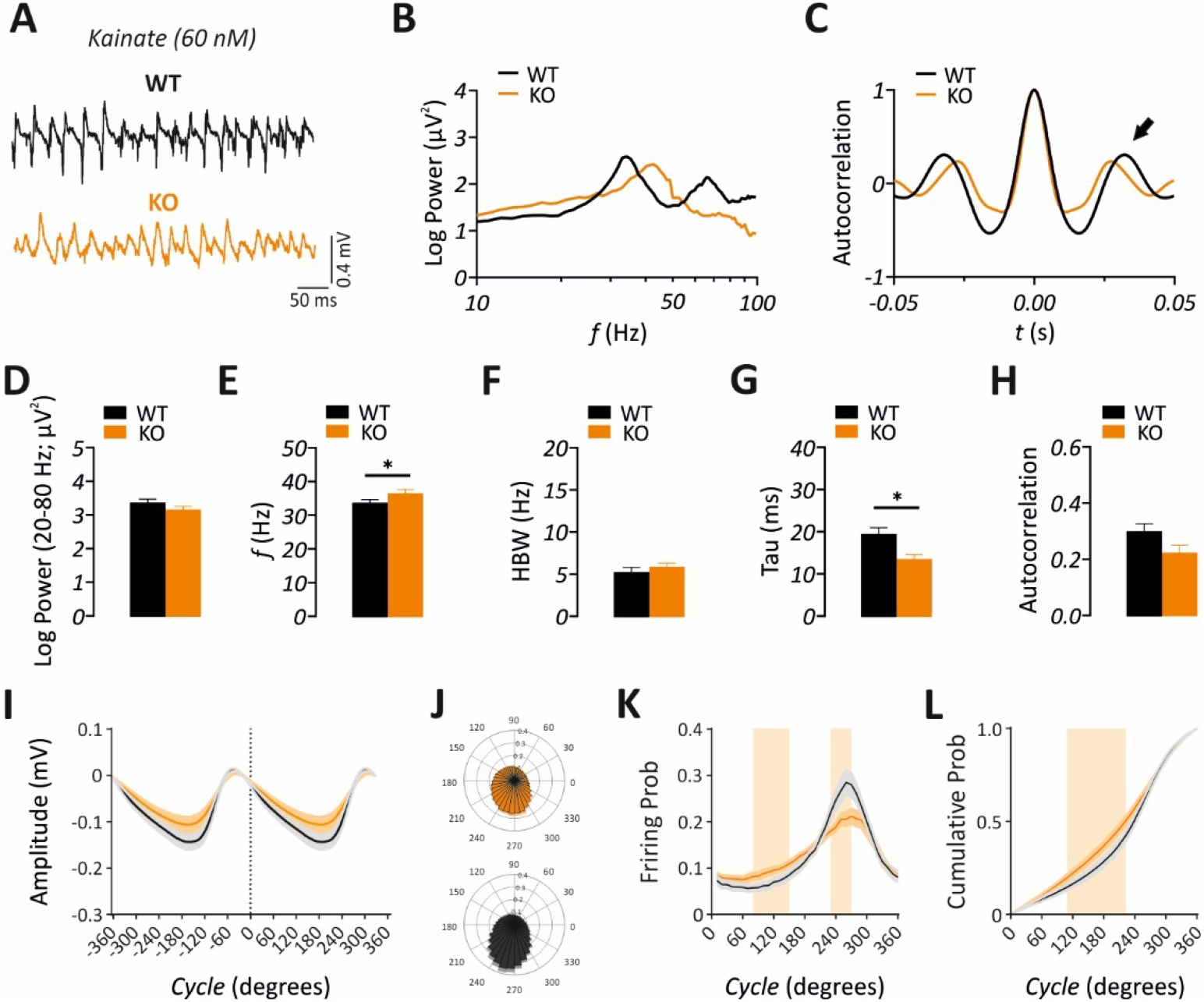
Kainate-induced gamma oscillations are reduced in their synchrony in the hippocampal CA3 of FMR1 KO mice *in vitro*. **(A)** Sample trace of gamma oscillations induced via bath-application of kainate (KA, 60 nM). Note the increased frequency and distorted synchrony of gamma oscillations in the FMR1 KO in comparison to the WT. **(B)** Corresponding power spectra of gamma oscillations showing a shift in the gamma peak oscillation frequency (*f*). **(C)** Corresponding autocorrelogram showing a slight reduction in the 2^nd^ positive peak value (indicated by the arrow) in the FMR1 KO. Summary plots illustrating a significant increase in the **(E)**the gamma frequency (WT: n = 31 / N = 7, KO: n = 30 / N = 7) and a significant decrease in **(G)**time constant (Tau) of the decaying exponential fit to the peaks of the autocorrelation (WT: n = 29 / N = 7, KO: n = 23 / N = 7), without strong alterations in **(D)** gamma power (20-80 Hz) (WT: n = 31 / N = 7, KO: n = 30 / N = 7), **(F)** half band width (HBW) (WT: n = 29 / N = 7, KO: n = 23 / N = 7) and **(H)** autocorrelation value (2^nd^ peak value of the autocorrelogram) (WT: n = 29 / N = 7, KO: n = 23 / N = 7). Evaluation of multi-unit activity in datasets (WT: n = 27 / N = 6, KO: n = 25 / N = 6) is shown in **I-L**. **(I)** Average gamma cycle. Note that the cycle’s amplitude is plotted against degrees. **(J)** Polar plots illustrating the probability of unit activity in FMR1 KO (top) and WT (bottom) mice relative to the cycle shown in **I**. **(K)** Unit firing probability reveals different activation regimes. Units in WT are significantly more active in the late phase, whereas units in FMR1 KO mice have a significantly higher firing probability in the early phase. **(L)** Unit activation curves reveal more uniform recruitment of units in FMR1 KO mice. Statistical comparison was performed using Student’s two-tailed t-test for **D, F, G** and Mann-Whitney U test for **C** and **E**. In **K** and **L** statistical comparison was done with multiple t-test comparisons. Colored patches indicate ranges in which datasets differed significantly. Data are given as mean ± SEM. *p < 0.05.

### 3.4. Spontaneous SW-R incidence is reduced in the hippocampal CA3 of FMR1 KO mice

Under reduced subcortical neuromodulation, the same hippocampal circuits can generate SW-R activity (Buzsáki, 2015). Thus, we assessed different parameters of SW-R in the hippocampal CA3 and CA1 subregions of FMR1 KO and WT mice (Fig. A-B). We found that SW-R incidence in both CA3 and CA1 areas were decreased in FMR1 KO mice compared to WT (Fig. 5C; CA3: Student’s two-tailed t-test; T(95) = 2.389, p < 0.05; CA1: Mann-Whitney U test, p < 0.05). However, we did not find any differences in other SW-R properties including SW area (Fig. 5D; CA3: Mann-Whitney U test, p = 0.4989; CA1: Mann-Whitney U test, p = 0.9898), ripple frequency (Fig. 5E; CA3: Mann-Whitney U test, p = 0.7313; CA1: Mann-Whitney U test, p = 0.5425) and ripple amplitude (Fig. 5F; CA3: Mann-Whitney U test, p = 0.6776; CA1: Mann-Whitney U test, p = 0.2080). These data indicate that SW rate in the ventral-to-mid hippocampus *in vitro* is overall reduced in the FMR1 KO mice.

**Fig. 5.**
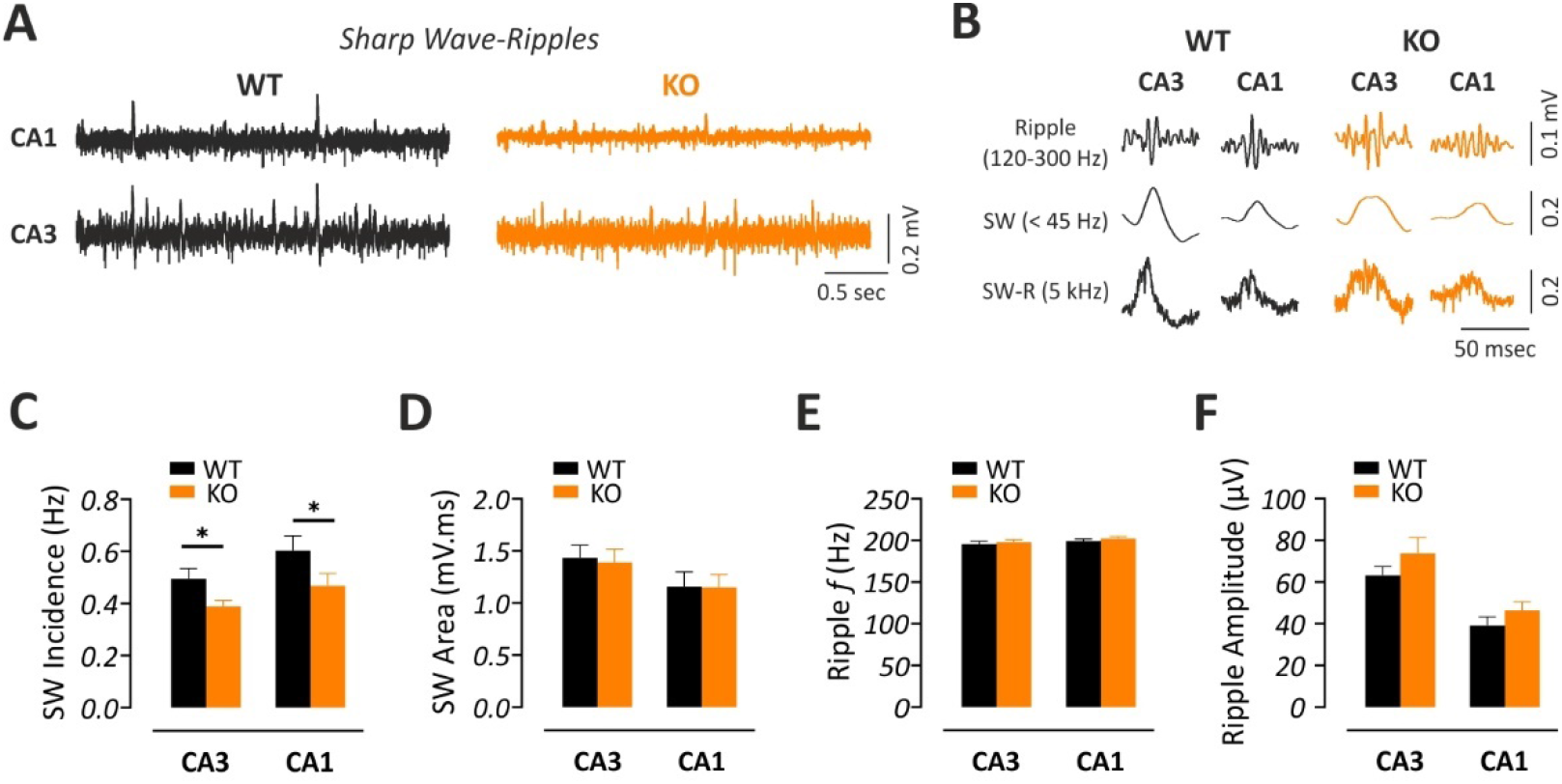
Decreased SW-R incidence in the hippocampal CA3 and CA1 of FMR1 KO mice *in vitro*. **(A)** Sample traces of spontaneous sharp wave-ripples (SW-R) in the CA3 and CA1 of WT and FMR1 KO mice, illustrating a decreased number of SW-R events. **(B)** Filtered SW-R traces showing the ripple component (band-pass filtered signal (120-300 Hz)) and the sharp wave component (low-pass filtered signal (<45 Hz)). Summary graphs showing **(C)** decreased incidence in the FMR1 KO compared to the WT, in both CA3 and CA1 areas with no changes in **(D)** SW area, **(E)** ripple frequency (*f*) and **(F)** ripple amplitude (CA3: WT, n = 43 / N = 11, KO, n = 54 / N = 14; CA1: WT, n = 38 / N = 11, KO, n = 49 / N = 14). Statistical comparison was performed using Student’s two-tailed t-test for **C** (CA3) and Mann-Whitney U test for **C** (CA1), **D**, **E** and **F**. Data are presented as mean ± SEM. *p < 0.05.

## 4. Discussion

In the current study, we took advantage of the well-established *in vitro* slice models for the analysis of gamma oscillations in a mouse model of FXS. We used three experimental protocols for the induction of gamma oscillations *in vitro* which involve pharmacological activation of either mAChR, mGluR or KAR (Fig. 1). Aberrant signaling mediated by activation of these receptors has been linked to the pathogenesis of FXS (Dölen et al., 2007; Michalon et al., 2012; Qiu et al., 2018; Thomson et al., 2017; Veeraragavan et al., 2011; Volk et al., 2007). Thus, this approach allowed us to assess potential contribution of these signaling pathways to the generation of hypersynchronous gamma oscillations in the FMR1 KO mouse model. Strikingly, we found that activation of either mAChR or mGluR leads to a profound increase in the power of gamma oscillations in the hippocampus of FMR1 KO. On the other hand, pharmacological activation of KAR leads to rather desynchronized gamma oscillations without altering the power of gamma oscillations suggesting a potential loss of KAR function in the FMR1 KO mouse. These changes were associated with altered cellular firing patterns during gamma oscillations induced by either CCh or KA. We further identified a mild reduction in the incidence of hippocampal SW-R events, LFP patterns which are linked to memory consolidation and appear during reduced subcortical neuromodulation (Buzsáki, 2015; Çalışkan and Stork, 2019). Together, our data suggest that aberrant activation of mAChR, mGluR or KAR may directly contribute to the gamma oscillopathy and associated behavioral abnormalities observed in FXS.

The most striking finding of the current study is the abnormal increase in the amplitude of cholinergic gamma oscillations in the hippocampal CA3 of FMR1 KO mouse (Fig. 2). Indeed, increased cholinergic tonus is associated with the emergence of LFP gamma oscillations in the hippocampus *in vivo* and *in vitro* (Caliskan et al., 2015; Fisahn et al., 1998; Vandecasteele et al., 2014). However, ACh metabolism appears to be normal in male FMR1 KO mice suggesting that the increased gamma oscillation power observed in FXS is not due to an overall increase in the availability of ACh (Scremin et al., 2015). Of note, generation of CCh-induced gamma oscillations depends on the activation of M1-AChR (Fisahn et al., 2002, 1998). Accordingly, FMR1 KO mouse model shows a significant increase in the expression of M1-AChR (Thomson et al., 2017) and an exaggerated CCh-induced hippocampal LTD mediated by M1-AChR is evident in the FMR1 KO mouse (Volk et al., 2007). These findings suggest that aberrant increase in the hippocampal gamma power of FXS might be due to elevated expression of M1-AChR and abnormal activation of downstream signaling pathways.

Similar to cholinergic gamma oscillations, we found a strong enhancement of mGluR1/5 agonist DHPG-induced gamma oscillations in the hippocampal CA3 of FMR1 KO mice (Fig. 3). Our results conform with previous studies (Pálhalmi et al., 2004) demonstrating a higher gamma peak frequency of DHPG-induced gamma oscillations in comparison to the other pharmacologically-induced gamma oscillations. These findings also align well with the enhanced mGluR signaling and mGluR-dependent LTD reported numerous times in the hippocampal CA1 of the FMR1 KO mice (Dölen et al., 2007; Dölen and Bear, 2008; Ferrante et al., 2021; Huber et al., 2002; Zhang et al., 2009). Of note, antagonism of mGluR appears to have no effect on both CCh-induced LTD and gamma oscillations *in vitro* (Fisahn et al., 1998; Pálhalmi et al., 2004; Volk et al., 2007). Interestingly, activation of either mGluR or M1-AChR leads to the enhancement of protein-synthesis dependent LTD via a common mechanism that is under the translational control of FMRP (Volk et al., 2007). Thus, synergistic actions of aberrant mGluR and M1-AChR signaling likely contribute to the enhanced gamma oscillations in the FMR1 KO mice.

Notably, KA-induced gamma oscillations were differently affected in the FMR1 KO mice (Fig. 4). We noted a general alteration in the oscillation parameters that indicate reduced synchronization of KA-induced gamma oscillations in the FMR1 KO mice. Previous pharmacological experiments demonstrate that KA-induced gamma oscillations do not require activation of either mAChR or mGluR suggesting that the potential reduction in the KAR function is independent of mAChR- or mGluR-mediated signaling mechanisms in the FMR1 KO mice (Fisahn et al., 2004). This finding aligns with the reduced synaptic expression and function of KAR in the cortex of the FMR1 KO mice (Qiu et al., 2018). To the best of our knowledge, no other studies directly investigated a potential change in the KAR function in the hippocampus of the FMR1 KO mice. Thus, KAR-dependent synaptic function and its molecular underpinnings in the hippocampus of the FMR1 KO mice need to be addressed in future studies.

CA3 LFP gamma oscillations represent compound synaptic potentials recorded extracellularly and their generation requires activity of both pyramidal cells and GABAergic interneurons (Hájos and Paulsen, 2009). Theoretical and experimental work indicate that distinct *in vitro* network oscillations require different levels of inhibitory and excitatory fast synaptic transmission. Blockade of GABA_A_R-mediated fast inhibitory transmission eliminates gamma oscillatory activity in all *in vitro* models measured in the current study (Fisahn et al., 2004, 1998; Pálhalmi et al., 2004). Drugs that prolong the fast inhibitory synaptic currents, such as zolpidem and pentobarbital, reduce the gamma peak frequency of KA- and DHPG-induced gamma oscillations (Fisahn et al., 2004; Pálhalmi et al., 2004), whereas this frequency shift is not observed for CCh-induced gamma oscillations (Pálhalmi et al., 2004). On the other hand, blockade of AMPAR-mediated fast excitatory synaptic transmission leads to complete elimination of CCh- and DHPG-induced gamma oscillations (Fisahn et al., 1998; Pálhalmi et al., 2004) without any major effect on KA-induced gamma oscillations (Fisahn et al., 2004). Collectively, with regards to the mouse model of FXS, these findings have several implications: (1) *in vitro* gamma oscillations (CCh- and DHPG-induced) which depend on an intact AMPAR-driven excitatory synaptic transmission are elevated, (2) inhibition-driven KA-induced gamma oscillations are reduced in their synchrony and show an increase in gamma peak frequency, (3) excitation-inhibition balance during gamma oscillations may underlie the distinct changes in the properties of gamma network activity induced by CCh, DHPG or KA in the hippocampal CA3 of FMR1 KO mice. Accordingly, hippocampal CA3 neurons of the FMR1 KO mice appear to be hyperexcitable and gamma-range stimulation of CA3 fibers leads to an aberrant augmentation of the excitatory neurotransmission at the CA3-CA1 synapses (Deng et al., 2019, 2011). Several studies also provide evidence for an altered excitation-inhibition balance in the FMR1 KO mice (Contractor et al., 2015; Kang et al., 2017; Sabanov et al., 2017). Indeed, disturbances in the hippocampal tonic and phasic inhibition together with lower expression of α2-subunit / δ-subunit containing GABA_A_R and R1a-containing GABA_B_R have been demonstrated (Kang et al., 2017; Sabanov et al., 2017). Hence, in future studies, it would be interesting to further explore a possible alteration in the excitation-inhibition balance during ongoing oscillations at the single cell level using e.g., patch clamp electrophysiology.

Parvalbumin-positive (PV+) inhibitory interneurons are indispensable for generation and maintenance of gamma oscillations both *in vivo* and *in vitro* (Buzsáki and Wang, 2012; Hájos and Paulsen, 2009). PV+ interneuron dysfunction and reduction in their number have been reported in the FMR1 KO mice (Lee et al., 2019; Lovelace et al., 2020; Steullet et al., 2017; Wen et al., 2018) and in postmortem neocortical tissue of autistic individuals (Hashemi et al., 2017). Paradoxically, deficient PV+ neuron function can lead to increased broadband gamma power, however, with a desynchronized cellular activity to the ongoing LFP oscillations (Guyon et al., 2021). Indeed, despite the overall increase in the pyramidal cell firing (Boone et al., 2018), such desynchronization of cellular activity within the CA1 network and to the ongoing LFP oscillations have been demonstrated in the FMR1 KO mice (Arbab et al., 2018; Talbot et al., 2018). In line with these findings, our analysis of extracellular unit firing in a gamma-cycle dependent manner revealed an aberrant cellular firing pattern during gamma oscillations induced by CCh or KA. We identified an aberrant increase in unit firing, most probably representing the action potentials generated by principal pyramidal cells, during the late stages of gamma cycles induced by CCh. On the other hand, a reduction in cellular firing was evident, when gamma oscillations were induced by KA. Thus, increased CCh-induced gamma oscillations might reflect such an aberrant increase in cellular firing leading to an enhanced broadband gamma power as observed *in vivo* (Arbab et al., 2018; Guyon et al., 2021; Talbot et al., 2018). On the other hand, reduced synchronization of gamma oscillations induced by KA might be associated with the desynchronization of underlying cellular activities (reduced unit firing) together with a potentially reduced KAR function in the hippocampus as reported in the cortex of the FMR1 KO mice (Qiu et al., 2018). Interestingly, we could not identify such an altered unit firing during DHPG-induced gamma oscillations. The increase in gamma power observed during DHPG-induced gamma oscillations in the FMR1 KO mice was rather small in comparison to CCh-induced gamma oscillations. In addition, our analysis of extracellular unit firing might not detect changes in subcellular activities (e.g. synaptic potentials or “spikelets”) which can also influence gamma oscillations strength. Hence, future studies combining single cell and extracellular electrophysiology will help resolving cellular correlates of increased DHPG-induced gamma oscillations in the FMR1 KO mice.

Our results also align with the findings of recent *in vivo* studies investigating the oscillatory changes in the hippocampus of FMR1 KO mice demonstrating a prominent dominance of CA3-driven slow gamma oscillations (20-50 Hz) in the CA1 region (Arbab et al., 2018; Boone et al., 2018; Dvorak et al., 2018; Radwan et al., 2016). In line, our recordings from the CA3 region of the hippocampus confirm a profound increase in the gamma oscillations induced by CCh- and DHPG with peak gamma frequencies ranging between 20-to-50 Hz (Fig. 2 and Fig. 3). Dominance of CA3-driven slow gamma oscillations was associated with aberrant memory encoding and retrieval, eventually leading to deficits in cognitive flexibility (Dvorak et al., 2018; Radwan et al., 2016; Talbot et al., 2018). These independent observations support the use of *in vitro* gamma oscillations as robust behaviorally-relevant physiological read-outs.

Under low cholinergic tonus, for example during slow wave sleep, hippocampal LFP activity is dominated by SW-R (Buzsaki, 1989; Buzsáki et al., 1992). These LFP oscillations have been linked to the replay of memory-related cellular activity, thus, supporting memory consolidation (Girardeau et al., 2009; Wilson and McNaughton, 1994). In slice preparations including the transverse sections of ventral-to-mid portion of the hippocampus, spontaneous SW-R activity also emerges and can be used as proxy to those that occur *in vivo* (Çalışkan et al., 2016; Mizunuma et al., 2014; Norimoto et al., 2018). We found an overall reduction in the incidence of hippocampal SW-R *in vitro* (Fig. 5). As aforementioned for gamma oscillations, generation and incidence of SW-R activity depends on functionally-intact PV+ interneurons (Ellender et al., 2010; Schlingloff et al., 2014). Thus, our results fit well with the potentially reduced PV+ interneuron function in the FMR1 KO mice leading to reduced SW-R incidence. To the best of our knowledge, only one recent study reports changes in the SW-R recorded in the dorsal hippocampus of the FMR1 KO mice demonstrating a reduced SW-R duration and ripple frequency without any alteration in SW-R incidence *in vivo* (Boone et al., 2018). Of note, major part of our recordings were performed from the ventral portion of the hippocampus which substantially differs functionally and anatomically from its dorsal counterpart (Fanselow and Dong, 2010). Thus, the discrepancies in the findings of the previous and current study might be due to different experimental settings (*in vivo* vs. *in vitro*) and hippocampal sections (dorsal vs. ventral) used in these two studies and merits further investigation. Interestingly, increased incidence of SW-R in the ventral hippocampus has been associated with an increased contextual fear memory consolidation (Çalışkan et al., 2016). Hence, a reduced SW-R incidence found in the current study fits well with the previously demonstrated contextual fear memory deficit in the FMR1 KO mice (Ding et al., 2014).

Taken together, the current study demonstrates that activation of either mAChR or mGluR is sufficient to mimic the aberrant increase in the gamma oscillations observed in the hippocampus of the FMR1 KO mice *in vivo*. It should be noted that the increase in gamma oscillation power has also been observed in numerous cortical areas including the auditory, somatosensory, frontal and temporal cortex of the FMR1 KO mice (Jonak et al., 2020; Lovelace et al., 2020, 2018; Pirbhoy et al., 2020; Wen et al., 2019; Wong et al., 2020). Whether these oscillatory alterations can be mimicked using pharmacologically-induced gamma oscillations in cortical slice preparations (Steullet et al., 2014) needs to be investigated in future. This will warrant the robust testing of potential pharmaceuticals in the treatment of neuronal hyperexcitability in FXS. It is important to note that human EEG studies have reported a similar enhancement of gamma power in individuals with FXS (Ethridge et al., 2019, 2017, 2016; Wang et al., 2017; Wilkinson and Nelson, 2021). Thus, utilization of gamma oscillation power as a mesoscopic biomarker may pave the way in the diagnosis of neurodevelopmental disorders such as FXS and facilitate target identification for the treatment of neuronal hyperexcitability in FXS.

## Acknowledgements

We are grateful to A. Koffi von Hoff, F. Blitz and S. Stork for excellent technical assistance and to A. Bohnstedt and D. Al-Chackmakchie for excellent animal care. This work was supported by the Center for Behavioural Brain Sciences - CBBS promoted by Europäische Fonds für regionale Entwicklung - EFRE (ZS/2016/04/78113) and CBBS - ScienceCampus funded by the Leibniz Association (SAS-2015-LIN-LWC) to GC. EP is a PhD student of ESF graduate school ABINEP (Funded by the federal state Saxony-Anhalt and the European Structural and Investment Funds (ESF, 2014-2020), ZS/2016/08/80645).

## References

Albrecht, A., Çalişkan, G., Oitzl, M.S., Heinemann, U., Stork, O., 2013. Long-lasting increase of corticosterone after fear memory reactivation: Anxiolytic effects and network activity modulation in the ventral hippocampus. Neuropsychopharmacology 38. https://doi.org/10.1038/npp.2012.192

Aloisi, E., Le Corf, K., Dupuis, J., Zhang, P., Ginger, M., Labrousse, V., Spatuzza, M., Georg Haberl, M., Costa, L., Shigemoto, R., Tappe-Theodor, A., Drago, F., Vincenzo Piazza, P., Mulle, C., Groc, L., Ciranna, L., Catania, M.V., Frick, A., 2017. Altered surface mGluR5 dynamics provoke synaptic NMDAR dysfunction and cognitive defects in Fmr1 knockout mice. Nat. Commun. 8. https://doi.org/10.1038/s41467-017-01191-2

Arbab, T., Battaglia, F.P., Pennartz, C.M.A., Bosman, C.A., 2018. Abnormal hippocampal theta and gamma hypersynchrony produces network and spike timing disturbances in the Fmr1-KO mouse model of Fragile X syndrome. Neurobiol. Dis. 114, 65–73. https://doi.org/10.1016/j.nbd.2018.02.011

Bakker, C.E., Verheij, C., Willemsen, R., van der Helm, R., Oerlemans, F., Vermey, M., Bygrave, A., Hoogeveen, A.T., Oostra, B.A., Reyniers, E., De Boule, K., D’Hooge, R., Cras, P., van Velzen, D., Nagels, G., Martin, J.J., De Deyn, P.P., Darby, J.K., Willems, P.J., 1994. Fmr1 knockout mice: A model to study fragile X mental retardation. Cell 78, 23–33. https://doi.org/10.1016/0092-8674(94)90569-X

Bartos, M., Vida, I., Jonas, P., 2007. Synaptic mechanisms of synchronized gamma oscillations in inhibitory interneuron networks. Nat. Rev. Neurosci. 8, 45–56. https://doi.org/10.1038/nrn2044

Bassell, G.J., Warren, S.T., 2008. Fragile X Syndrome: Loss of Local mRNA Regulation Alters Synaptic Development and Function. Neuron 60, 201–214. https://doi.org/10.1016/j.neuron.2008.10.004

Bear, M.F., Huber, K.M., Warren, S.T., 2004. The mGluR theory of fragile X mental retardation. Trends Neurosci. 27, 370–377. https://doi.org/10.1016/j.tins.2004.04.009

Boone, C.E., Davoudi, H., Harrold, J.B., Foster, D.J., 2018. Abnormal Sleep Architecture and Hippocampal Circuit Dysfunction in a Mouse Model of Fragile X Syndrome. Neuroscience 384, 275–289. https://doi.org/10.1016/j.neuroscience.2018.05.012

Butler, J.L., Paulsen, O., 2015. Hippocampal network oscillations — recent insights from in vitro experiments. Curr. Opin. Neurobiol. 31, 40–44. https://doi.org/10.1016/j.conb.2014.07.025

Buzsaki, G., 1989. Two-stage model of memory trace formation: A role for “noisy” brain states. Neuroscience 31, 551–570.

Buzsáki, G., 2015. Hippocampal sharp wave-ripple: A cognitive biomarker for episodic memory and planning. Hippocampus 25, 1073–1188. https://doi.org/10.1002/hipo.22488

Buzsaki, G., Draguhn, A., 2004. Neuronal Oscillations in Cortical Networks. Science (80-.). 304, 1926–1929. https://doi.org/10.1126/science.1099745

Buzsáki, G., Horváth, Z., Urioste, R., Hetke, J., Wise, K., 1992. High-frequency network oscillation in the hippocampus. Science 256, 1025–1027. https://doi.org/10.1126/science.1589772

Buzsáki, Wang, X.-J., 2012. Mechanisms of Gamma Oscillations. Annu. Rev. Neurosci. 203–225. https://doi.org/10.1146/annurev-neuro-062111-150444

Caliskan, G., Schulz, S.B., Gruber, D., Behr, J., Heinemann, U., Gerevich, Z., 2015. Corticosterone and corticotropin-releasing factor acutely facilitate gamma oscillations in the hippocampus in vitro. Eur. J. Neurosci. 41, 31–44. https://doi.org/10.1111/ejn.12750

Çalışkan, G., Müller, I., Semtner, M., Winkelmann, A., Raza, A.S., Hollnagel, J.O., Rösler, A., Heinemann, U., Stork, O., Meier, J.C., 2016. Identification of Parvalbumin Interneurons as Cellular Substrate of Fear Memory Persistence. Cereb. Cortex 26, 2325–2340. https://doi.org/10.1093/cercor/bhw001

Çalışkan, G., Stork, O., 2019. Hippocampal network oscillations at the interplay between innate anxiety and learned fear. Psychopharmacology (Berl). 236, 321–338. https://doi.org/10.1007/s00213-018-5109-z

Çalışkan, G., Stork, O., 2018. Hippocampal network oscillations as mediators of behavioural metaplasticity: Insights from emotional learning. Neurobiol. Learn. Mem. 154, 37–53. https://doi.org/10.1016/j.nlm.2018.02.022

Contractor, A., Klyachko, V.A., Portera-Cailliau, C., 2015. Altered Neuronal and Circuit Excitability in Fragile X Syndrome. Neuron 87, 699–715. https://doi.org/10.1016/j.neuron.2015.06.017

Deng, P.Y., Carlin, D., Oh, Y.M., Myrick, L.K., Warren, S.T., Cavalli, V., Klyachko, V.A., 2019. Voltage-independent SK-channel dysfunction causes neuronal hyperexcitability in the hippocampus of Fmr1 knock-out mice. J. Neurosci. 39, 28–43. https://doi.org/10.1523/JNEUROSCI.1593-18.2018

Deng, P.Y., Sojka, D., Klyachko, V.A., 2011. Abnormal presynaptic short-term plasticity and information processing in a mouse model of fragile X syndrome. J. Neurosci. 31, 10971–10982. https://doi.org/10.1523/JNEUROSCI.2021-11.2011

Ding, Q., Sethna, F., Wang, H., 2014. Behavioral analysis of male and female Fmr1 knockout mice on C57BL/6 background. Behav. Brain Res. 271, 72–78. https://doi.org/10.1186/s12967-016-1105-4

Dölen, G., Bear, M.F., 2008. Role for metabotropic glutamate receptor 5 (mGluR5) in the pathogenesis of fragile X syndrome. J. Physiol. 586, 1503–1508. https://doi.org/10.1113/jphysiol.2008.150722

Dölen, G., Osterweil, E., Rao, B.S.S., Smith, G.B., Auerbach, B.D., Chattarji, S., Bear, M.F., 2007. Correction of Fragile X Syndrome in Mice. Neuron 56, 955–962. https://doi.org/10.1016/j.neuron.2007.12.001

Dvorak, D., Radwan, B., Sparks, F.T., Talbot, Z.N., Fenton, A.A., 2018. Control of recollection by slow gamma dominating mid-frequency gamma in hippocampus CA1. PLoS Biol. 16, 1–27. https://doi.org/10.1371/journal.pbio.2003354

Ellender, T.J., Nissen, W., Colgin, L.L., Mann, E.O., Paulsen, O., 2010. Priming of hippocampal population bursts by individual perisomatic-targeting interneurons. J. Neurosci. 30, 5979–91. https://doi.org/10.1523/JNEUROSCI.3962-09.2010

Ethridge, L.E., De Stefano, L.A., Schmitt, L.M., Woodruff, N.E., Brown, K.L., Tran, M., Wang, J., Pedapati, E. V., Erickson, C.A., Sweeney, J.A., 2019. Auditory EEG Biomarkers in Fragile X Syndrome: Clinical Relevance. Front. Integr. Neurosci. 13, 1–16. https://doi.org/10.3389/fnint.2019.00060

Ethridge, L.E., White, S.P., Mosconi, M.W., Wang, J., Byerly, M.J., Sweeney, J.A., 2016. Reduced habituation of auditory evoked potentials indicate cortical hyper-excitability in fragile X syndrome. Transl. Psychiatry 6. https://doi.org/10.1038/tp.2016.48

Ethridge, L.E., White, S.P., Mosconi, M.W., Wang, J., Pedapati, E. V., Erickson, C.A., Byerly, M.J., Sweeney, J.A., 2017. Neural synchronization deficits linked to cortical hyper-excitability and auditory hypersensitivity in fragile X syndrome. Mol. Autism 8, 1–11. https://doi.org/10.1186/s13229-017-0140-1

Fanselow, M.S., Dong, H.W., 2010. Are the Dorsal and Ventral Hippocampus Functionally Distinct Structures? Neuron 65, 7–19. https://doi.org/10.1016/j.neuron.2009.11.031

Ferrante, A., Boussadia, Z., Borreca, A., Mallozzi, C., Pedini, G., Pacini, L., Pezzola, A., Armida, M., Vincenzi, F., Varani, K., Bagni, C., Popoli, P., Martire, A., 2021. Adenosine A2A receptor inhibition reduces synaptic and cognitive hippocampal alterations in Fmr1 KO mice. Transl. Psychiatry 11. https://doi.org/10.1038/s41398-021-01238-5

Fisahn, A., Contractor, A., Traub, R.D., Buhl, E.H., Heinemann, S.F., McBain, C.J., 2004. Distinct Roles for the Kainate Receptor Subunits GluR5 and GluR6 in Kainate-Induced Hippocampal Gamma Oscillations. J. Neurosci. 24, 9658–9668. https://doi.org/10.1523/JNEUROSCI.2973-04.2004

Fisahn, A., Pike, F.G., Buhl, E.H., Paulsen, O., 1998. Cholinergic induction of network oscillations at 40 Hz in the hippocampus in vitro. Nature 394, 186–189. https://doi.org/10.1038/28179

Fisahn, A., Yamada, M., Duttaroy, A., Gan, J.W., Deng, C.X., McBain, C.J., Wess, J., 2002. Muscarinic induction of hippocampal gamma oscillations requires coupling of the M1 receptor to two mixed cation currents. Neuron 33, 615–624. https://doi.org/10.1016/S0896-6273(02)00587-1

Gillies, M.J., Traub, R.D., LeBeau, F.E.N., Davies, C.H., Gloveli, T., Buhl, E.H., Whittington, M.A., 2002. A model of atropine-resistant theta oscillations in rat hippocampal area CA1. J. Physiol. 543, 779– 793. https://doi.org/10.1113/jphysiol.2002.024588

Girardeau, G., Benchenane, K., Wiener, S.I., Buzsáki, G., Zugaro, M.B., 2009. Selective suppression of hippocampal ripples impairs spatial memory. Nat. Neurosci. 12, 1222–1223. https://doi.org/10.1038/nn.2384

Guyon, N., Zacharias, L.R., de Oliveira, E.F., Kim, H., Leite, J.P., Lopes-Aguiar, C., Carlén, M., 2021. Network asynchrony underlying increased broadband gamma power. J. Neurosci. JN-RM-2250-20. https://doi.org/10.1523/JNEUROSCI.2250-20.2021

Hagerman, R.J., Berry-Kravis, E., Hazlett, H.C., Bailey, D.B., Moine, H., Kooy, R.F., Tassone, F., Gantois, I., Sonenberg, N., Mandel, J.L., Hagerman, P.J., 2017. Fragile X syndrome. Nat. Rev. Dis. Prim. 3, 17065. https://doi.org/10.1038/nrdp.2017.65

Hájos, N., Pálhalini, J., Mann, E.O., Nèmeth, B., Paulsen, O., Freund, T.F., 2004. Spike timing of distinct types of GABAergic interneuron during hippocampal gamma oscillations in vitro. J. Neurosci. 24, 9127–9137. https://doi.org/10.1523/JNEUROSCI.2113-04.2004

Hájos, N., Paulsen, O., 2009. Network mechanisms of gamma oscillations in the CA3 region of the hippocampus. Neural Networks 22, 1113–1119. https://doi.org/10.1016/j.neunet.2009.07.024

Hashemi, E., Ariza, J., Rogers, H., Noctor, S.C., Martínez-Cerdeño, V., 2017. The Number of Parvalbumin-Expressing Interneurons Is Decreased in the Medial Prefrontal Cortex in Autism. Cereb. Cortex 27, 1931–1943. https://doi.org/10.1093/cercor/bhw021

Hollnagel, J.O., Elzoheiry, S., Gorgas, K., Kins, S., Beretta, C.A., Kirsch, J., Kuhse, J., Kann, O., Kiss, E., 2019. Early alterations in hippocampal perisomatic GABAergic synapses and network oscillations in a mouse model of Alzheimer’s disease amyloidosis. PLoS One 14, 1–23. https://doi.org/10.1371/journal.pone.0209228

Huber, K.M., Gallagher, S.M., Warren, S.T., Bear, M.F., 2002. Altered synaptic plasticity in a mouse model of fragile X mental retardation. Proc. Natl. Acad. Sci. U. S. A. 99, 7746–7750. https://doi.org/10.1073/pnas.122205699

Jonak, C.R., Lovelace, J.W., Ethell, I.M., Razak, K.A., Binder, D.K., 2020. Multielectrode array analysis of EEG biomarkers in a mouse model of Fragile X Syndrome. Neurobiol. Dis. 138, 104794. https://doi.org/10.1016/j.nbd.2020.104794

Kang, J.Y., Chadchankar, J., Vien, T.N., Mighdoll, M.I., Hyde, T.M., Mather, R.J., Deeb, T.Z., Pangalos, M.N., Brandon, N.J., Dunlop, J., Moss, S.J., 2017. Deficits in the activity of presynaptic-aminobutyric acid type B receptors contribute to altered neuronal excitability in fragile X syndrome. J. Biol. Chem. 292, 6621–6632. https://doi.org/10.1074/jbc.M116.772541

Klein, A.S., Donoso, J.R., Kempter, R., Schmitz, D., Beed, P., 2016. Early Cortical Changes in Gamma Oscillations in Alzheimer’s Disease. Front. Syst. Neurosci. 10, 83. https://doi.org/10.3389/fnsys.2016.00083

Le Duigou, C., Simonnet, J., Teleñczuk, M.T., Fricker, D., Miles, R., 2014. Recurrent synapses and circuits in the CA3 region of the hippocampus: An associative network. Front. Cell. Neurosci. 7, 1–13. https://doi.org/10.3389/fncel.2013.00262

Lee, F.H.F., Lai, T.K.Y., Su, P., Liu, F., 2019. Altered cortical Cytoarchitecture in the Fmr1 knockout mouse. Mol. Brain 12, 1–12. https://doi.org/10.1186/s13041-019-0478-8

Lovelace, J.W., Ethell, I.M., Binder, D.K., Razak, K.A., 2018. Translation-relevant EEG phenotypes in a mouse model of Fragile X Syndrome. Neurobiol. Dis. 115, 39–48.

Lovelace, J.W., Rais, M., Palacios, A.R., Shuai, X.S., Bishay, S., Popa, O., Pirbhoy, P.S., Binder, D.K., Nelson, D.L., Ethell, I.M., Razak, K.A., 2020. Deletion of Fmr1 from Forebrain Excitatory Neurons Triggers Abnormal Cellular, EEG, and Behavioral Phenotypes in the Auditory Cortex of a Mouse Model of Fragile X Syndrome. Cereb. Cortex 30, 969–988. https://doi.org/10.1093/cercor/bhz141

Lu, C.B., Jefferys, J.G.R., Toescu, E.C., Vreugdenhil, M., 2011. In vitro hippocampal gamma oscillation power as an index of in vivo CA3 gamma oscillation strength and spatial reference memory. Neurobiol. Learn. Mem. 95, 221–230. https://doi.org/10.1016/j.nlm.2010.11.008

Maier, N., Nimmrich, V., Draguhn, A., 2003. Cellular and network mechanisms underlying spontaneous sharp wave-ripple complexes in mouse hippocampal slices. J. Physiol. 550, 873–887. https://doi.org/10.1113/jphysiol.2003.044602

Michalon, A., Sidorov, M., Ballard, T.M., Ozmen, L., Spooren, W., Wettstein, J.G., Jaeschke, G., Bear, M.F., Lindemann, L., 2012. Chronic Pharmacological mGlu5 Inhibition Corrects Fragile X in Adult Mice. Neuron 74, 49–56. https://doi.org/10.1016/j.neuron.2012.03.009

Mizunuma, M., Norimoto, H., Tao, K., Egawa, T., Hanaoka, K., Sakaguchi, T., Hioki, H., Kaneko, T., Yamaguchi, S., Nagano, T., Matsuki, N., Ikegaya, Y., 2014. Unbalanced excitability underlies offline reactivation of behaviorally activated neurons. Nat. Neurosci. 17, 503–5. https://doi.org/10.1038/nn.3674

Norimoto, H., Makino, K., Gao, M., Shikano, Y., Okamoto, K., Ishikawa, T., Sasaki, T., Hioki, H., Fujisawa, S., Ikegaya, Y., 2018. Hippocampal ripples down-regulate synapses. Science (80-.). 359, 1524–1527. https://doi.org/10.1126/science.aao0702

Pálhalmi, J., Paulsen, O., Freund, T.F., Hájos, N., 2004. Distinct properties of carbachol- and DHPG-induced network oscillations in hippocampal slices. Neuropharmacology 47, 381–389. https://doi.org/10.1016/j.neuropharm.2004.04.010

Pilpel, Y., Kolleker, A., Berberich, S., Ginger, M., Frick, A., Mientjes, E., Oostra, B.A., Seeburg, P.H., 2009. Synaptic ionotropic glutamate receptors and plasticity are developmentally altered in the CA1 field of Fmr1 knockout mice. J. Physiol. 587, 787–804. https://doi.org/10.1113/jphysiol.2008.160929

Pirbhoy, P.S., Rais, M., Lovelace, J.W., Woodard, W., Razak, K.A., Binder, D.K., Ethell, I.M., 2020. Acute pharmacological inhibition of matrix metalloproteinase-9 activity during development restores perineuronal net formation and normalizes auditory processing in Fmr1 KO mice. J. Neurochem. 155, 538–558. https://doi.org/10.1111/jnc.15037

Qiu, S., Wu, Y., Lv, X., Li, X., Zhuo, M., Koga, K., 2018. Reduced synaptic function of Kainate receptors in the insular cortex of Fmr1 Knock-out mice. Mol. Brain 11, 1–11. https://doi.org/10.1186/s13041-018-0396-1

Radwan, B., Dvorak, D., Fenton, A.A., 2016. Impaired cognitive discrimination and discoordination of coupled theta-gamma oscillations in Fmr1 knockout mice. Neurobiol. Dis. 88, 125–138. https://doi.org/10.1016/j.nbd.2016.01.003

Sabanov, V., Braat, S., D’Andrea, L., Willemsen, R., Zeidler, S., Rooms, L., Bagni, C., Kooy, R.F., Balschun, D., 2017. Impaired GABAergic inhibition in the hippocampus of Fmr1 knockout mice. Neuropharmacology 116, 71–81. https://doi.org/10.1016/j.neuropharm.2016.12.010

Salcedo-Arellano, M.J., Dufour, B., McLennan, Y., Martinez-Cerdeno, V., Hagerman, R., 2020. Fragile X syndrome and associated disorders: Clinical aspects and pathology. Neurobiol. Dis. 136, 104740. https://doi.org/10.1016/j.nbd.2020.104740

Santos, A.R., Kanellopoulos, A.K., Bagni, C., 2014. Learning and behavioral deficits associated with the absence of the fragile X mental retardation protein: What a fly and mouse model can teach us. Learn. Mem. 21, 543–555. https://doi.org/10.1101/lm.035956.114

Schlingloff, D., Kali, S., Freund, T.F., Hajos, N., Gulyas, A.I., 2014. Mechanisms of Sharp Wave Initiation and Ripple Generation. J. Neurosci. 34, 11385–11398. https://doi.org/10.1523/JNEUROSCI.0867-14.2014

Scremin, O.U., Roch, M., Norman, K.M., Djazayeri, S., Liu, Y.Y., 2015. Brain acetylcholine and choline concentrations and dynamics in a murine model of the Fragile X syndrome: Age, sex and region-specific changes. Neuroscience 301, 520–528. https://doi.org/10.1016/j.neuroscience.2015.06.036

Sinclair, D., Oranje, B., Razak, K.A., Siegel, S.J., Schmid, S., 2017. Sensory processing in autism spectrum disorders and Fragile X syndrome—From the clinic to animal models. Neurosci. Biobehav. Rev. 76, 235–253. https://doi.org/10.1016/j.neubiorev.2016.05.029

Steullet, P., Cabungcal, J.-H., Cuénod, M., Do, K.Q., 2014. Fast oscillatory activity in the anterior cingulate cortex: dopaminergic modulation and effect of perineuronal net loss. Front. Cell. Neurosci. 8, 244. https://doi.org/10.3389/fncel.2014.00244

Steullet, P., Cabungcal, J.H., Coyle, J., Didriksen, M., Gill, K., Grace, A.A., Hensch, T.K., Lamantia, A.S., Lindemann, L., Maynard, T.M., Meyer, U., Morishita, H., O’Donnell, sP., Puhl, M., Cuenod, M., Do, K.Q., 2017. Oxidative stress-driven parvalbumin interneuron impairment as a common mechanism in models of schizophrenia. Mol. Psychiatry 22, 936–943. https://doi.org/10.1038/mp.2017.47

Talbot, Z.N., Sparks, F.T., Dvorak, D., Curran, B.M., Alarcon, J.M., Fenton, A.A., 2018. Normal CA1 Place Fields but Discoordinated Network Discharge in a Fmr1-Null Mouse Model of Fragile X Syndrome. Neuron 97, 684–697.e4. https://doi.org/10.1016/j.neuron.2017.12.043

Thomson, S.R., Seo, S.S., Barnes, S.A., Louros, S.R., Muscas, M., Dando, O., Kirby, C., Wyllie, D.J.A., Hardingham, G.E., Kind, P.C., Osterweil, E.K., 2017. Cell-Type-Specific Translation Profiling Reveals a Novel Strategy for Treating Fragile X Syndrome. Neuron 95, 550–563.e5. https://doi.org/10.1016/j.neuron.2017.07.013

Vandecasteele, M., Varga, V., Berényi, A., Papp, E., Barthó, P., Venance, L., Freund, T.F., Buzsáki, G., 2014. Optogenetic activation of septal cholinergic neurons suppresses sharp wave ripples and enhances theta oscillations in the hippocampus. Proc. Natl. Acad. Sci. U. S. A. 111, 13535–40. https://doi.org/10.1073/pnas.1411233111

Veeraragavan, S., Bui, N., Perkins, J.R., Yuva-Paylor, L.A., Carpenter, R.L., Paylor, R., 2011. Modulation of behavioral phenotypes by a muscarinic M1 antagonist in a mouse model of fragile X syndrome. Psychopharmacology (Berl). 217, 143–151. https://doi.org/10.1007/s00213-011-2276-6

Volk, L.J., Pfeiffer, B.E., Gibson, J.R., Huber, K.M., 2007. Multiple Gq-coupled receptors converge on a common protein synthesis-dependent long-term depression that is affected in fragile X syndrome mental retardation. J. Neurosci. 27, 11624–11634. https://doi.org/10.1523/JNEUROSCI.2266-07.2007

Wang, J., Ethridge, L.E., Mosconi, M.W., White, S.P., Binder, D.K., Pedapati, E. V., Erickson, C.A., Byerly, M.J., Sweeney, J.A., 2017. A resting EEG study of neocortical hyperexcitability and altered functional connectivity in fragile X syndrome Refining translational treatment development in fragile X syndrome. J. Neurodev. Disord. 9, 1–12. https://doi.org/10.1186/s11689-017-9191-z

Wen, T.H., Afroz, S., Reinhard, S.M., Palacios, A.R., Tapia, K., Binder, D.K., Razak, K.A., Ethell, I.M., 2018. Genetic Reduction of Matrix Metalloproteinase-9 Promotes Formation of Perineuronal Nets Around Parvalbumin-Expressing Interneurons and Normalizes Auditory Cortex Responses in Developing Fmr1 Knock-Out Mice. Cereb. Cortex 28, 3951–3964. https://doi.org/10.1093/cercor/bhx258

Wen, T.H., Lovelace, J.W., Ethell, I.M., Binder, D.K., Razak, K.A., 2019. Developmental Changes in EEG Phenotypes in a Mouse Model of Fragile X Syndrome. Neuroscience 398, 126–143. https://doi.org/10.1016/j.neuroscience.2018.11.047

Wilkinson, C.L., Nelson, C.A., 2021. Increased aperiodic gamma power in young boys with Fragile X is associated with better language ability. Mol. Autism 12. https://doi.org/10.1101/2020.10.08.20209536

Wilson, M.A., McNaughton, B.L., 1994. Reactivation of hippocampal ensemble memories during sleep. Science (80-.). 265, 676–679. https://doi.org/10.1126/science.8036517

Wójtowicz, A.M., Van Boom, L. Den, Chakrabarty, A., Maggio, N., Haq, R.U., Behrens, C.J., Heinemann, U., 2009. Monoamines block kainate- And carbachol-induced. γ-oscillations but augment stimulus-induced γ-oscillations in rat hippocampus in vitro. Hippocampus 19, 273–288. https://doi.org/10.1002/hipo.20508

Wong, H., Hooper, A.W.M., Niibori, Y., Lee, S.J., Hategan, L.A., Zhang, L., Karumuthil-Melethil, S., Till, S.M., Kind, P.C., Danos, O., Bruder, J.T., Hampson, D.R., 2020. Sexually dimorphic patterns in electroencephalography power spectrum and autism-related behaviors in a rat model of fragile X syndrome. Neurobiol. Dis. 146. https://doi.org/10.1016/j.nbd.2020.105118

Zemankovics, R., Veres, J.M., Oren, I., Hajos, N., 2013. Feedforward Inhibition Underlies the Propagation of Cholinergically Induced Gamma Oscillations from Hippocampal CA3 to CA1. J. Neurosci. 33, 12337–12351. https://doi.org/10.1523/JNEUROSCI.3680-12.2013

Zhang, J., Hou, L., Klann, E., Nelson, D.L., 2009. Altered hippocampal synaptic plasticity in the Fmr1 gene family knockout mouse models. J. Neurophysiol. 101, 2572–2580. https://doi.org/10.1152/jn.90558.2008

